# Does the leaf economic spectrum hold within plant functional types? A Bayesian multivariate trait meta-analysis

**DOI:** 10.1101/475038

**Authors:** Alexey N. Shiklomanov, Elizabeth M. Cowdery, Michael Bahn, Chaeho Byun, Steven Jansen, Koen Kramer, Vanessa Minden, Ülo Niinemets, Yusuke Onoda, Nadejda A. Soudzilovskaia, Michael C. Dietze

**Author notes:** Corresponding author; Mail: 5825 University Research Ct., Office 3533, College Park, MD 20740.

## Abstract

We investigated whether global leaf economic relationships are also present within plant functional types (PFTs), and the extent to which this hierarchical structure can be used to constrain trait estimates. We developed a hierarchical multivariate Bayesian model that assumes separate means and covariance structures within and across PFTs and fit this model to seven leaf traits from the TRY database related to leaf morphology, biochemistry, and photosynthetic metabolism. Trait correlations were generally consistent in direction within and across PFTs, and consistent with predictions of the leaf economic spectrum. However, correlation strength varied substantially across PFTs indicating that leaf economic relationships within PFTs are often confounded by the unique physiology of certain plant types or environmental conditions in certain biomes. Leveraging covariance in multivariate models reduced uncertainties in mean trait estimates, particularly for undersampled trait-PFT combinations. However, additional constraint from the across-PFT hierarchy was limited.

**Data accessibility:** The R code and ancillary data for running these analyses is publicly available online via the Open Science Framework at https://osf.io/w8y73/. The TRY data request used for this analysis has been archived at http://try-db.org, and can be retrieved by providing the TRY data request ID (#1584). Alternatively, the exact preformatted data used in this analysis are available on request to Alexey Shiklomanov.

## 1 Introduction

The use of functional groups with common characteristics is widespread in ecological studies (Naeem Wright 2003) and is essential to many ecosystem models (Lavorel *et al.* 1997; Wullschleger *et al.* 2014). However, ecologists have long recognized the importance of individual variability and stochasticity in shaping ecosystems (Gleason 1939; Bolnick *et al.* 2011; Rosindell *et al.* 2011; Clark 2016) and the benefits of more finely-resolved representation of functional diversity are supported by an increasing body of ecological literature (Mayfield *et al.* 2006; McMahon *et al.* 2011; Van Bodegom *et al.* 2011; Reichstein *et al.* 2014; Violle *et al.* 2014; Medlyn *et al.* 2015; Moran *et al.* 2015). Plant functional traits can be used to link directly measurable features of individuals to their fitness within an ecosystem, and to whole-ecosystem function (Violle *et al.* 2007). Recent trait syntheses have revealed that plant functional diversity is constrained by allometries and trade-offs between ecological strategies (Wright *et al.* 2004; Kattge *et al.* 2011; Díaz *et al.* 2015; Kleyer & Minden 2015). One such constraint is the “leaf economic spectrum", which defines a trade-off between productive but short-lived leaves versus less productive but recalcitrant and long-lived leaves (Wright *et al.* 2004; Shipley *et al.* 2006; Reich 2014; Díaz *et al.* 2015). Leaf economic traits are well-correlated with plant productivity (Shipley *et al.* 2005; Niinemets 2016; Wu *et al.* 2016b), litter decomposition rates (Bakker *et al.* 2010; Hobbie 2015), community composition (Burns 2004; Cavender-Bares *et al.* 2004), and ecosystem function (Diaz *et al.* 2004; Musavi *et al.* 2015). The position of species along the leaf economic spectrum is influenced by climate and soil conditions (Wright *et al.* 2004, 2005; Cornwell & Ackerly 2009; Ordoñez *et al.* 2009; Wigley *et al.* 2016). As a result, relationships between leaf economic traits and climate have been used in ecosystem models for more finely-resolved variation in plant function (Sakschewski *et al.* 2015; Verheijen *et al.* 2015).

However, the use of global among-trait and trait-environment correlations, for both ecological inference and predictive modeling, has several important caveats. First, observed global correlations may not hold at smaller scales, such as sites, species, and individuals. Some studies suggest consistent correlations across scales (Wright *et al.* 2004; Albert *et al.* 2010a; Asner *et al.* 2014), whereas others show no or even opposite correlations (Albert *et al.* 2010b; Messier *et al.* 2010, 2017; Wright & Sutton-Grier 2012; Feng & Dietze 2013; Kichenin *et al.* 2013; Grubb *et al.* 2015; Wigley *et al.* 2016). Many mechanisms have been suggested for scale-dependence of trait relationships. Trade-offs may only apply when multiple competing strategies co-occur, and alternative processes can drive community assembly where strong environmental filters severely limit the range of feasible strategies (Grime & Pierce 2012; Rosado & Mattos 2017). Different selective pressures dominate at different scales, particularly within versus across species (Albert *et al.* 2010b; Messier *et al.* 2010; Kichenin *et al.* 2013), and different traits have different sensitivities to such pressures (Messier *et al.* 2016). Experimental evidence shows that species can alter different aspects of their leaf economy independently (Wright & Sutton-Grier 2012). Meanwhile, allocation patterns (and therefore investment strategies and trait relationships) vary across plant functional types (Ghimire *et al.* 2017). Second, among-trait correlations at any scale do not provide causal evidence for functional trade-offs (Messier *et al.* 2016), and ascribing excessive leverage to trait correlations can result in underestimation of functional diversity (Grubb 2015). Third, plants maintain their fitness through multiple independent strategies, corresponding to multiple mutually orthogonal axes of trait variability. For instance, changes in leaf economic traits can fail to predict changes in other aspects of plant function, such as hydraulics (Li *et al.* 2015), dispersal (Westoby *et al.* 2002), and overall plant carbon budget (Edwards *et al.* 2014). Finally, modeling ecosystem function based on trait correlations is sampling from the hypothetical space of potential species and communities that could have evolved, failing to account for real constraints such as the timescales required for adaptation and community assembly.

An alternative approach is to preserve existing PFT classifications (potentially with finer resolution, e.g. (Boulangeat *et al.* 2012)) while using statistical analyses to account for uncertainty and variability in the aggregated trait values. For example, the Predictive Ecosystem Analyzer (PEcAn, pecanproject.org), an ecosystem model-data informatics system, parameterizes PFTs using trait probability distributions from a Bayesian meta-analysis of trait data (Dietze *et al.* 2013; LeBauer *et al.* 2013). This approach explicitly separates the processes driving PFT-level differentiation from those driving finer-scale functional variability, and is useful for guiding future data collection and model refinement (Dietze *et al.* 2014). However, a univariate meta-analysis like PEcAn’s fails to account for trait correlations and therefore neglects useful knowledge about relationships across PFTs and between traits. At the other extreme, existing regional and global analyses (e.g. (Van Bodegom *et al.* 2011; Sakschewski *et al.* 2015)) ignore variability within PFTs, often resulting in macroecological, evolutionary, and competitive trade-offs across PFTs being used to drive acclimation and instantaneous responses within PFTs.

While the leaf economic spectrum has been investigated at the global scale, where it is robust, and at local scales, where deviations from it are common, it has received less attention at the intermediate scale of PFTs. Thus, this paper seeks to answer the following questions: First, does the leaf economic spectrum hold within vs. across PFTs? Second, can the leaf economic spectrum and similar covariance patterns be leveraged to constrain trait estimates, particularly under data limitation? The answers to these question have implications for both functional ecology and ecosystem modeling. To address these questions, we developed a hierarchical multivariate Bayesian model that explicitly accounts for across- and within-PFT variability in trait correlations. We then fit this model to a global trait database to estimate mean trait values and variance-covariance matrices for PFTs as defined in a major earth system model (Community Land Model, CLM, Oleson *et al.* 2013). We evaluate the ability of this model to reduce uncertainties in trait estimates and reproduce observed patterns of global trait variation compared to univariate models. Finally, we assess the scale dependence and generality of estimated trait covariances.

## 2 Materials and methods

### 2.1 Trait data

We obtained trait data from the TRY global database (Kattge *et al.* 2011) (see Appendix S1 in Supporting Information). We focused on seven foliar traits: longevity (months), specific leaf area (SLA, m^2^ kg^−1^), nitrogen content (*N*_*mass*_, mg N g^−1^ or *N*_*area*_, g m^−2^), phosphorus content (*P*_*mass*_, mg P g^−1^ or *P*_*area*_, g m^−2^), dark respiration at 25°C (*R*_*d*_,_*mass*_, pmol g^−1^ s^−1^, or *R*_*d,area*_, pmol m^−2^ s^−1^), _max_imum RuBisCO carboxylation rate at 25°C (*V*_*c*_,_*max*_,_*mass*_, pmol g^−1^ s^−1^, or *V*_*c*_,_*max*_,_*area*_, pmol m^−2^ s^−1^), and maximum electron transport rate at 25°C (J_*max,mass*_, μmol g^−1^ s^−1^, or J_*max,area*_, μmol m^−2^ s^−1^). For *Vc,*_*max*_, we only used values reported at 25°C. For Rd, we normalized the values to 25°C using reported leaf temperature values following (Atkin *et al.* 2015). For *J*_*max*_, we normalized the values to 25°C using reported leaf temperature values using the temperature response function described in (Kattge & Knorr 2007) (Equation 1 therein). To avoid issues with trait normalization, we performed analyses separately for both mass- and area-normalized traits (Lloyd *et al.* 2013; Osnas *et al.* 2013). We restricted our analysis to quality-controlled values from species with sufficient information for functional type classification (see Kattge *et al.* 2011). Following past studies (e.g. Wright *et al.* 2004; Onoda *et al.* 2011; Díaz *et al.* 2015), we logtransformed all trait values to correct for their strong right-skewness.

### 2.2 Plant functional types

We assigned each species a PFT following the scheme in the Community Land Model (CLM4.5, Oleson *et al.* 2013) (Tab. 1, Fig. 1). We obtained categorical data on growth form, leaf type, phenology, and photosynthetic pathway from TRY. Where species attributes disagreed between datasets, we assigned the most frequently observed attribute (e.g., if five datasets say “shrub” but only one says “tree", we would use “shrub”). Where species attributes were missing, we assigned attributes based on higher order phylogeny if possible (e.g., *Poaceae* family are grasses, *Larix* genus are deciduous needleleaved trees) or omitted the species if not. For biome specification, we matched geographic coordinates for each species to annual mean temperature (*AMT*, averaged 1970-2000) data from WorldClim-2 (Fick & Hijmans 2017), calculated the mean AMT for all sites where each species was observed, and then binned these species based on the following cutoffs: boreal/arctic *(AMT* ≤ 5°C), temperate (*AMT* ≤ 20°C), and tropical (*AMT* > 20°C).

**Figure 1:**
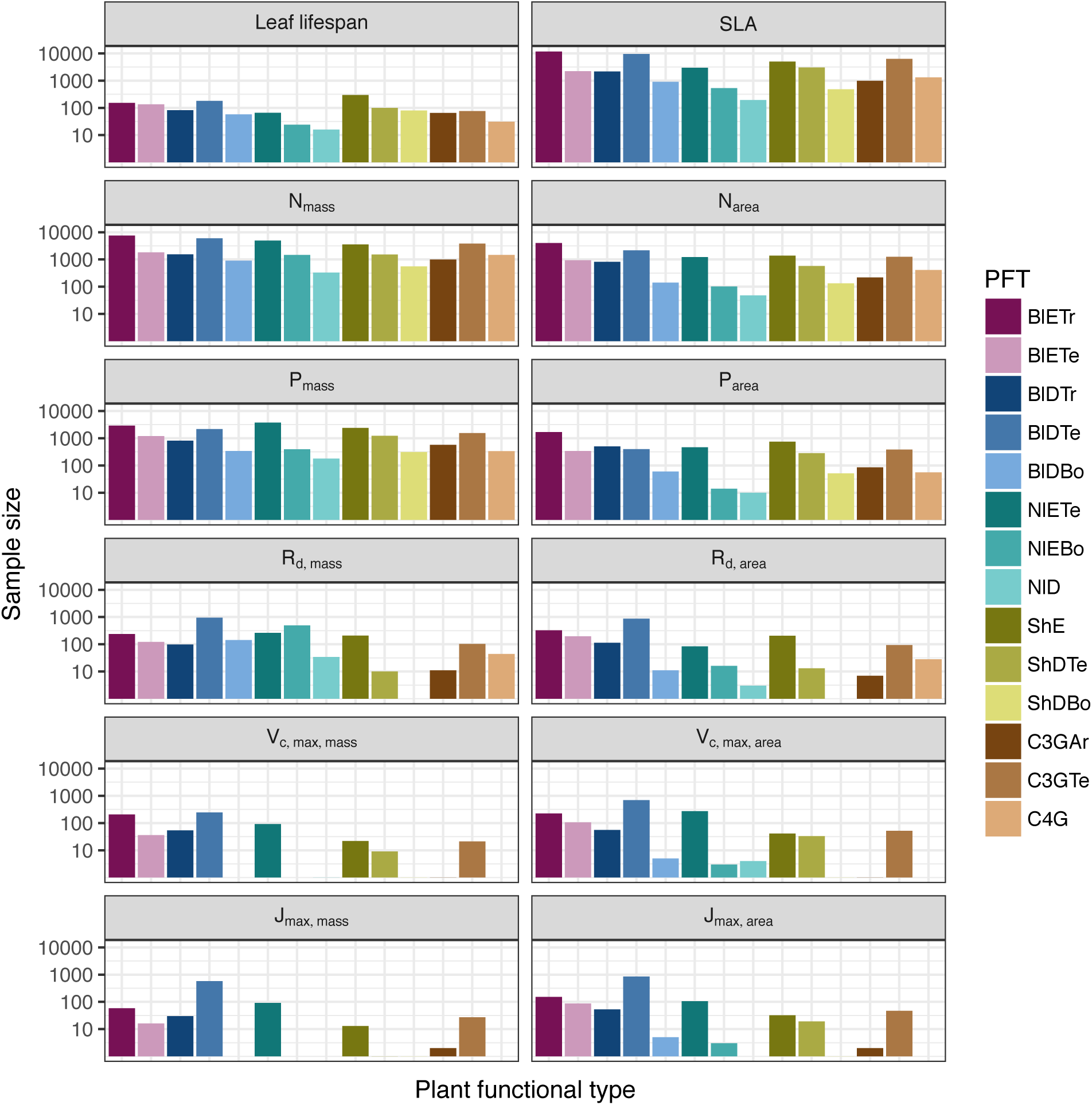
Sample sizes for each trait-PFT pair. *y* axis is scaled logarithmically.

### 2.3 Multivariate analysis

#### 2.3.1 Basic model description

We compared three models representing different levels of complexity. The simplest was the “univariate” model, in which each trait is independent. For an observation *x*_*i,t*_ of trait *t* and sample *i*:

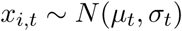

where *N* is the univariate Gaussian distribution with mean *μ*_*t*_ and standard deviation *σ*_*t*_ for trait *t*.

The second-simplest model was the “multivariate” model, in which traits are drawn from a single multivariate distribution. For observed trait vector x_i_ for sample *i*:

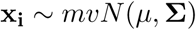

where *mvN* is the multivariate Gaussian distribution with mean vector *μ*and variance-covariance matrix Σ. We fit both of these models independently for each PFT and once for the entire dataset (i.e. one global PFT).

The most complex model was the “hierarchical” model, where traits are drawn from a PFTspecific multivariate distribution describing within-PFT variation, and whose mean vector is itself sampled from a global multivariate distribution describing variation across PFTs. For observed trait vector *x*_*i,p*_ for sample *i* belonging to PFT *p:*

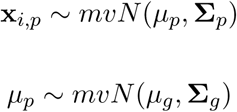

where *μ*_*p*_ and *Σ* _*p*_ are the mean vector and variance-covariance matrix describing variation within PFT p, and *μ* _*g*_ and *Σ*_*g*_ are the mean vector and variance-covariance matrix describing across-PFT (global) variation.

#### 2.3.2 Model implementation

We fit the above models using Gibbs sampling, which leverages conjugate prior relationships for efficient exploration of the sampling space. For priors on all multivariate mean vectors (μ), we used normal distributions:

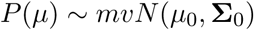

This leads to the following posterior distribution:

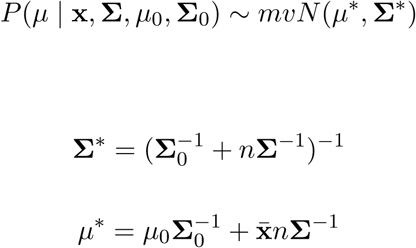

where 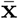 are the sample means of the data and *n* is the number of rows in the data.

For priors on all multivariate variance-covariance matrices, we used the Wishart distribution (*W*):

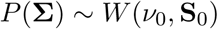

This leads to the following posterior distribution:

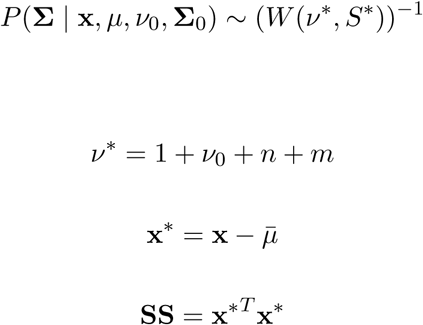

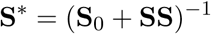

where *n* is the number of observations and *m* is the number of traits in data matrix x. For further details on this derivation, see (Gelman *et al.* 2003).

The multivariate nature of this sampling procedure makes it incapable of accommodating partially missing observations. Therefore, our algorithm included multiple imputation of partially missing data (Graham 2009; White *et al.* 2010). For a block of data x/ containing missing observations in columns m and present observations in columns p, missing values x/[m] are drawn randomly from a conditional multivariate normal distribution at each iteration of the sampling algorithm:

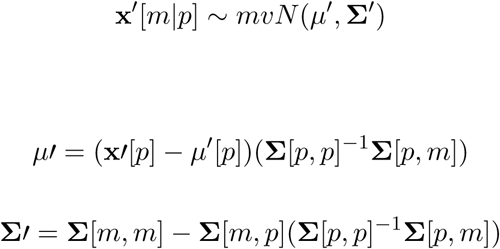

We emphasize that imputation of missing data is performed iteratively as parameters are being estimated, such that each set of imputed values is conditioned on the current sampled state of the parameters. This approach is distinct from single imputation, where data are imputed first in a separate step prior to parameter estimation (Graham 2009; White *et al.* 2010). A demonstration of this multiple imputation approach and how it is used to estimate trait covariance is provided Supporting Information Method S1.

For each model fit, we ran five sampling sequences, continuing sampling until the final result achieved convergence as determined by a Gelman-Rubin potential scale reduction statistic less than 1.1 (Gelman & Rubin 1992). We implemented this sampling algorithm in a publicly available R (R Core Team 2018) package (http://github.com/ashiklom/mvtraits).

#### 2.3.3 Analysis of results

To assess multivariate and hierarchical constraint on trait estimates, we compared the mean and 95% confidence intervals of trait estimates for each PFT from each model (Fig. 2, Tab. S1 and S2). For reference, we included the default parameter values of CLM 4.5 (Table Oleson *et al.* 2013) for SLA, *N*_*mass*_, *N*_*area*_, *V*_*c,max,mass*_, and *V*_*c,max,area*_ to Fig. 2. To convert CLM’s reported C:N ratio to *N*_*mass*_, we assumed a uniform leaf C fraction of 0.46. We then divided this calculated *N*_*mass*_ by the reported SLA to obtain *N*_*area*_. We calculated *V*_*c,max,mass*_ by multiplying the reported *Vc,*_*max,area*_ by the reported SLA.

**Figure 2:**
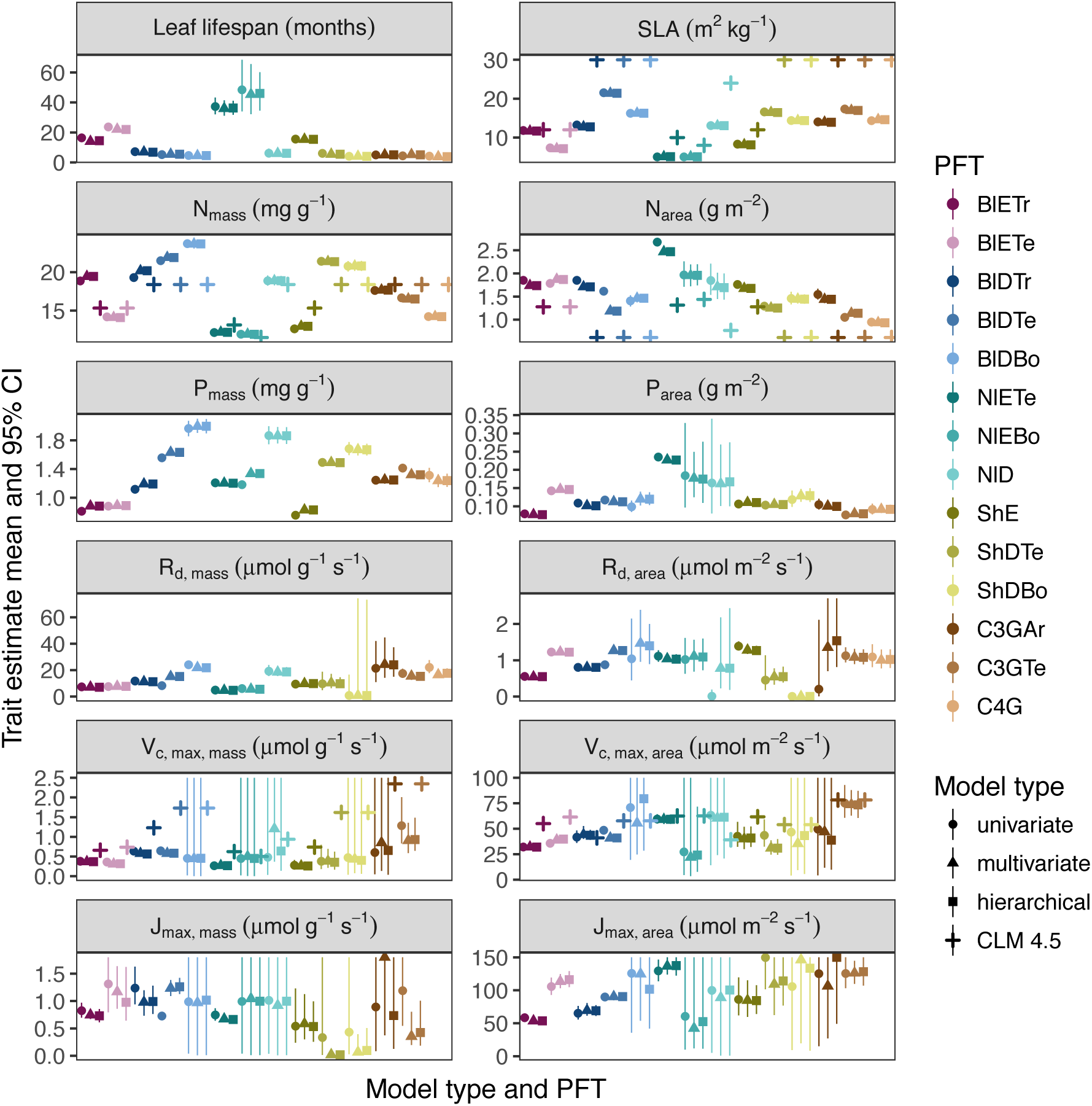
Mean and 95% confidence interval on best estimates of traits for each plant functional type from the univariate, multivariate, and hierarchical models. For leaf lifespan and SLA, results were not significantly different between the mass- and area-based models, so only results from the mass-based model are shown. For some PFT-trait combinations, where large error bars resulting from the relatively uninformative priors are substantially larger than the variability among means, the y axes are constrained to facilitate comparison.

To test whether multivariate and hierarchical models offer relatively more utility at smaller sample sizes, we calculated the relative uncertainty (*α*) as a function of the mean (*μ*) and upper (*q*_0.975_) and lower (*q*_0.025_) confidence limits of trait estimates.

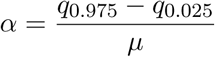

We then fit a log-linear least-squares regression relating relative uncertainty to sample size (*n*) for each model (univariate, multivariate, and hierarchical; Fig. 3).

**Figure 3:**
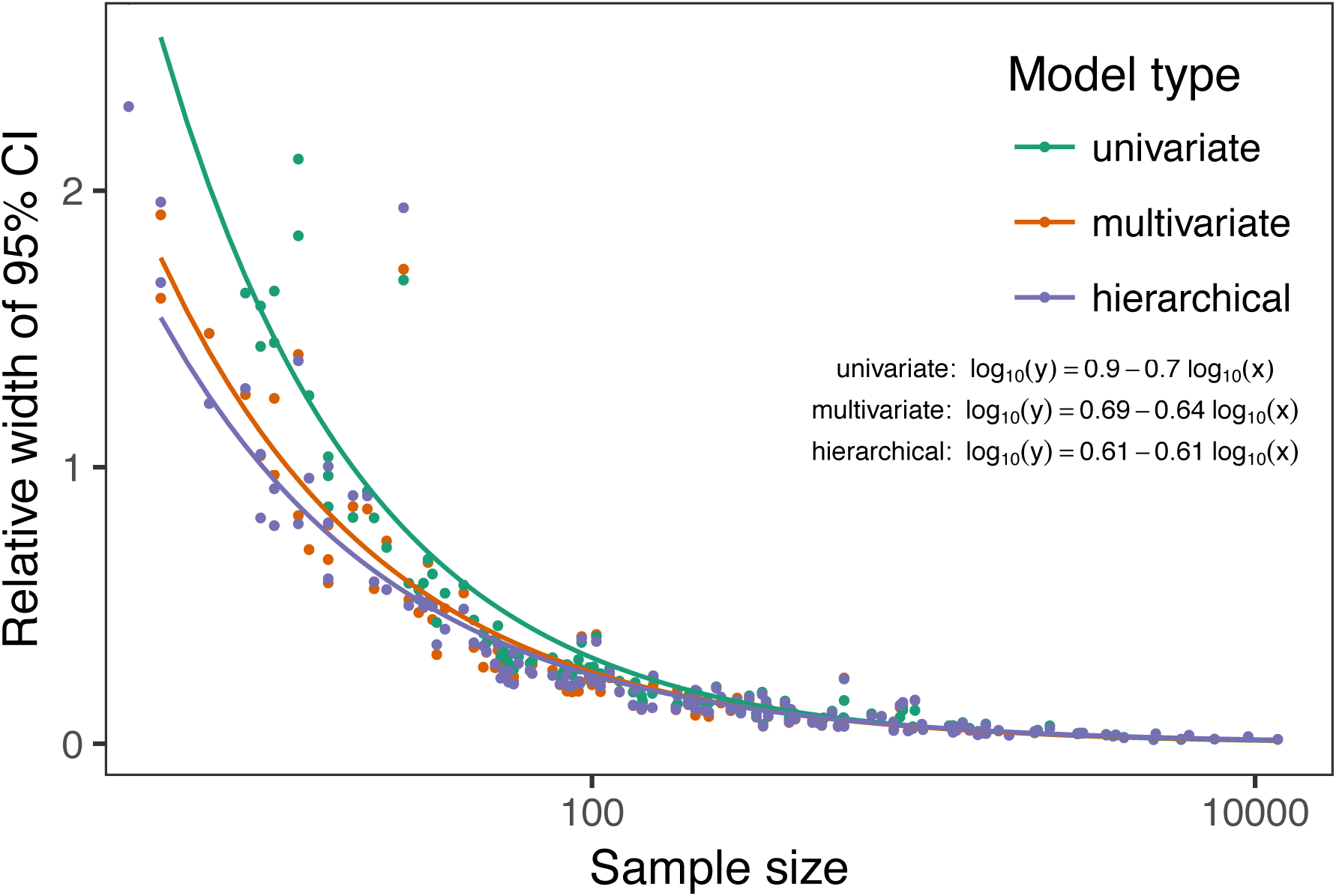
Relative uncertainty in PFT-level trait estimates as a function of sample size for each model type. Lines represent linear models (log(*y*) = *b*_*0*_ + *b*_*1*_ log(*x*)) fit independently for each model type. In general, differences in estimate uncertainty between the univariate and multivariate models were minimal at large sample sizes but increasingly important at low sample sizes. However, differences in estimate uncertainty between the multivariate and hierarchical models were consistently negligible.

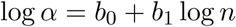

If all three models performed equally well at all sample sizes, their respective slope and intercept coefficients would be statistically indistinguishable. Meanwhile, models that perform better should have a lower intercept (*b*_0_), indicating lower overall uncertainty, and a lower slope (*b*_1_), indicating reduced sensitivity of uncertainty (*α*) to sample size (*n*).

To assess the consistency of within- and across-PFT trait trade-offs, we examined covariance estimates for each trait pair and, where these values were significantly different from zero (*p <* 0.05), we calculated the eigenvalues from the variance-covariance matrix for just that trait pair and plotted the corresponding dominant eigenvectors centered on the mean estimates (Fig. 4). This figure provides a visual representation of relative positions of PFTs in trait space and both the direction and extent of within-PFT trait covariance. It is analogous to conceptual figures describing hierarchical trait variability across environmental gradients as presented in (Cornwell & Ackerly 2009) and (Albert *et al.* 2010b). Due to the small number of points used to estimate across-PFT covariance in the hierarchical model, none of its across-PFT covariances were significantly different from zero (p < 0.05). Therefore, we compared within-PFT covariances from the hierarchical model against covariances from a single global multivariate model. Besides the consistency in the direction of trait covariance within vs. across PFTs, we also investigated the strength and predictive power of these covariances, represented by correlation coefficients (i.e. pairwise covariance normalized by each trait’s variance). We plotted the mean and 95% confidence interval of the pairwise trait correlation coefficients from the global multivariate model and PFT-specific estimates from the hierarchical model (Fig. 5).

**Figure 4:**
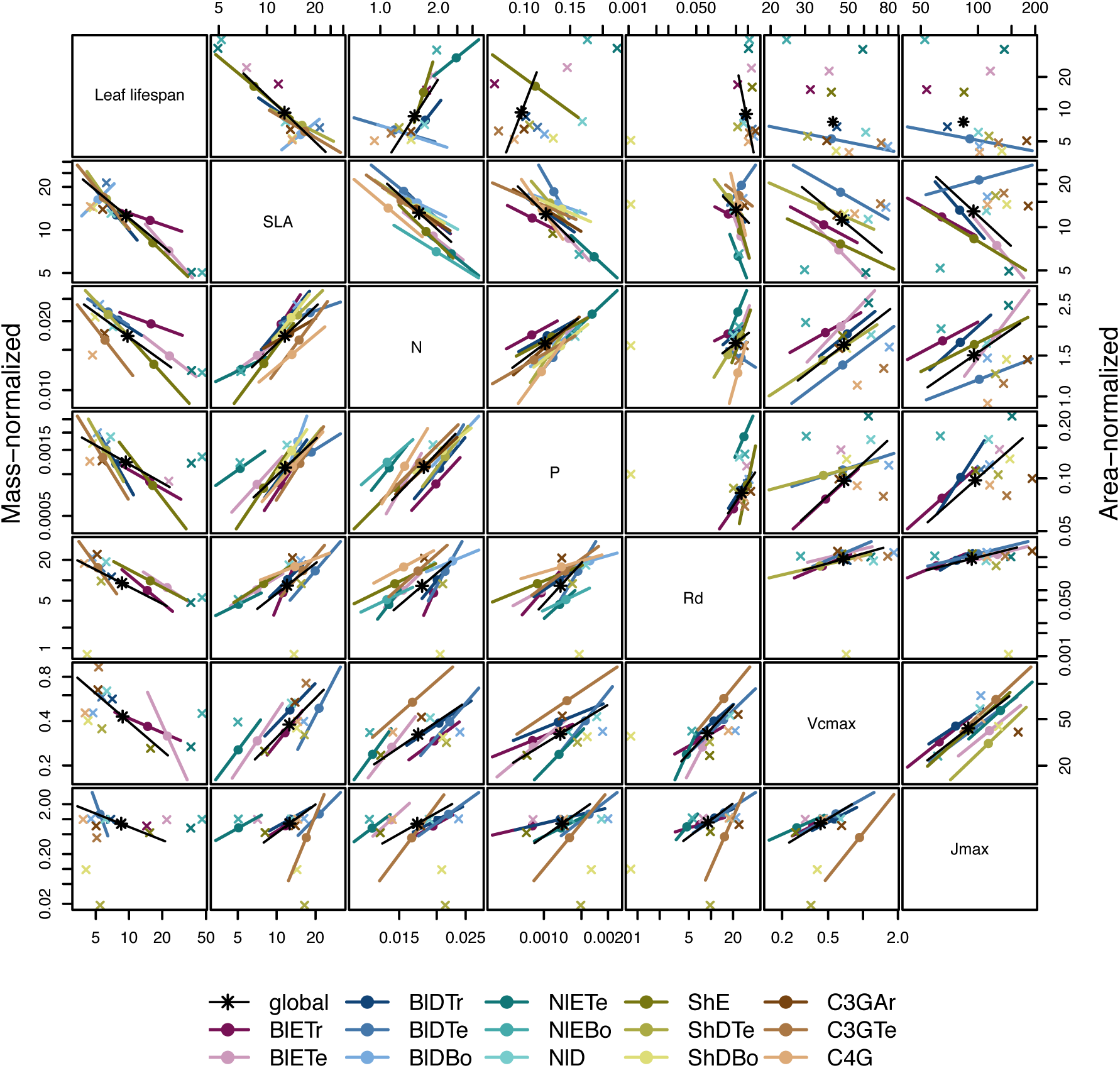
Pairwise trait mean and covariance estimates for all data pooled globally (black) and for each PFT (colored). Covariance estimates not significantly different from zero (*p <* 0.05) are indicated by x symbols at the mean estimate. *x* and *y* axes vary on a log scale, reflecting the fact that the model was fit using the base 10 log of all traits. With the exception of leaf lifespan, pairwise covariances are consistent in direction but vary somewhat in magnitude between PFTs, and when comparing PFT-level and global estimates. However, many pairwise covariances are not statistically significant, particularly (but not always) for undersampled traits and PFTs.

**Figure 5:**
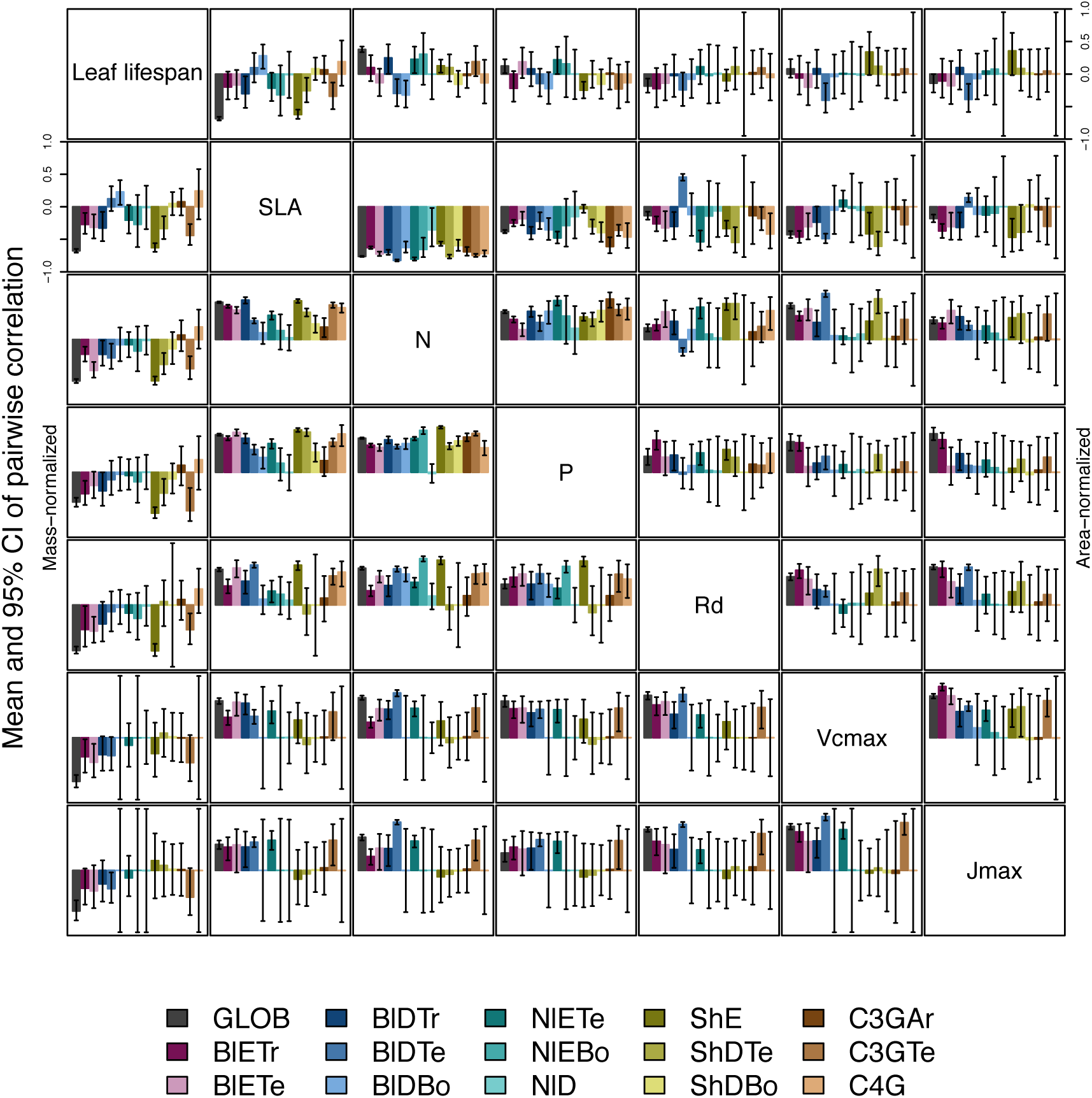
Mean and 95% CI on estimates of pairwise correlation coefficients for all data pooled globally (dark grey) and for each PFT (colored). For most PFT-trait pairs, correlations are mutually consistent in magnitude but vary in strength.

Correlation coefficients are sensitive to data sampling, particularly sample size and range (stronger correlations when data have more samples and larger range). To evaluate the contribution of data sampling to our correlation estimates, we plotted the each pairwise correlation coefficient squared against pairwise sample size and normalized data range (Fig. S1 and S2).

## 3 Results

### 3.1 Estimates of PFT-level means

In general, leaf trait estimates from the univariate, multivariate, and hierarchical models were similar (Fig. 2, Tab. S1 and S2). Where estimates differed between models, the largest differences were between the univariate and multivariate models, and additional constraint from the hierarchical model relative to PFT-specific multivariate models had a minimal effect on trait estimates. Significant differences in trait estimates between univariate and multivariate models occurred even for well-sampled traits, such as leaf nitrogen content.

Across-PFT patterns in SLA and *N*_*mass*_, *P*_*mass*_, and *Rd,*_*mass*_ were similar, with the highest values in temperate broadleaved deciduous PFTs and the lowest values in temperate evergreen PFTs. However, none of these patterns was universal to all four traits. For example, tropical evergreen trees had relatively high *N*_*mass*_ and average SLA and *R*_*d,mass*_, but among the lowest *P*_*mass*_. Similarly, compared to grass PFTs, temperate and boreal shrubs had similar SLA but higher *N*_*mass*_ and *P*_*mass*_. Patterns were different when these traits were normalized by _area_ instead of _mass_. For example, tropical broadleaved evergreen and needleleaf evergreen trees had relatively low *N*_*mass*_ and *P*_*mass*_ but relatively high *N*_*area*_ and *P*_*area*_, while the opposite was true of deciduous temperate trees and shrubs.

A key application of this study was to provide data-driven parameter estimates for Earth System Models. To this end, we compared our mean parameter estimates with corresponding default parameters in CLM 4.5 (Oleson *et al.* 2013) (Fig. 2). Our SLA estimates were significantly lower than CLM parameters for all PFTs except tropical broadleaved evergreen trees. Our *N*_*mass*_ estimates showed more across-PFT variability than CLM parameters, and only agreed with CLM for evergreen temperate trees, needleleaved trees, and C3 arctic grasses. Similarly to (Kattge *et al.* 2009), we found that CLM overestimates *V*_*c,max*_, both by mass and area.

We observed differences in the uncertainties of mean estimates with respect to sample size. High-latitude PFTs had large uncertainties relative to other PFTs, and the traits with the largest uncertainties were dark respiration, *V*_*c,max*_, and *J*_*max*_. For many of these trait-PFT combinations, the additional constraint from trait covariance provided by the multivariate and hierarchical models reduced error bars, making it possible to compare estimates against those of other PFTs. Our analysis of the relationship between sample size and trait uncertainty found that, compared to the univariate model, the multivariate model both reduced uncertainty overall (lower intercept) and reduced the sensitivity of uncertainty to sample size (lower slope) (Fig. 3). However, the additional benefit from the hierarchical model was small.

### 3.2 Trait correlation patterns across- and within-PFTs

For all traits except leaf lifespan, pairwise trait correlations were generally consistent in direction both globally and within each PFT (Fig. 4). Mass- and area-normalized traits were all positively correlated with each other and, respectively, positively and negatively correlated with SLA, both globally and within each PFT. Mass-based traits were generally positively correlated with leaf lifespan, but correlations of area-based traits with leaf lifespan were more variable. The *N*_*area*_ -leaf lifespan relationship was positive globally and for evergreen shrubs, tropical broadleaved deciduous trees, temperate needleleaved evergreen trees, but negative for temperate and boreal broadleaved deciduous trees and not significant for any other PFTs. Similarly, the correlation between *P*_*area*_ and leaf lifespan was positive globally but negative for evergreen shrubs and not significant for any other PFTs. The correlation between leaf lifespan and *R*_*d,area*_ was significant and negative globally, but was not significant within any PFTs. The only significant correlations of leaf lifespan with *V*_*c,max,area*_ and *J*_*max,area*_ were negative for temperate broadleaved deciduous trees.

Pairwise trait correlation strength varied depending on scale, PFT, and trait (Fig. 5). In some cases, this variability was driven by low sample sizes (Fig. 1, S1; Tab. S3, S4). For instance, needleleaved deciduous trees, the most undersampled PFT in our analysis, were often the only PFT for which a correlation was not statistically significant. Similarly, we had no observations of dark respirations for deciduous boreal shrubs, which explains why we found no significant correlations of dark respiration with any other trait for that PFT. However, the relationship between correlations strength and sample size was inconsistent (Fig. S1; Tab. S4). Every trait pair had at least one case (and often several cases) where a better-sampled PFT showed weaker correlations than PFTs with lower sample sizes, or where correlation strength varied significantly among PFTs with similar sample sizes (Fig. S1). Relationships between correlation strength and data range were even less consistent (Fig. S2). Therefore, we conclude that the variation in our correlation results can not be explained by sampling alone and captures some underlying ecophysiological differences between PFTs.

## 4 Discussion

### 4.1 Scale dependence of the leaf economic spectrum

The leaf economic spectrum is defined by a negative correlation between SLA and leaf lifespan, and a positive correlation of SLA with *N*_*mass*_, *P*_*mass*_, and photosynthesis and respiration rates (Wright et *al.* 2004). Our first objective was to investigate the extent to which these relationships hold within and across PFTs. Our results indicate that the leaf economic spectrum generally holds within PFTs, at least at the functional and phylogenetic resolution of current Earth System Models. Within PFTs, correlations between SLA, *N*_*mass*_, and *P*_*mass*_ were consistently positive, and correlations of these traits with leaf lifespan were generally negative (though, for many PFTs, correlations were not significantly different from zero). Although we did not include _max_imum photosynthesis rate (*A*_*max*_) ^*V*^*c,*_*max,mass*_ and *J*_*max,mass*_ generally exhibited the expected positive correlations with SLA and negative correlations with leaf lifespan, as did *R*_*d,mass*_, though many correlations were not significant.

While trait relationships within PFTs were consistent in direction, their strength was more variable. For example, correlations of SLA with *N*_*mass*_ and *P*_*mass*_ were weaker in needleleaved PFTs compared to broadleaved PFTs. Meanwhile, correlations of SLA with *N*_*area*_ were strongly negative for all PFTs (except the data-limited needleleaved deciduous trees), and especially so in temperate needleleaved species. Given that evergreen conifers have a relatively constant allocation of N to cell walls and RuBisCO (Onoda *et al.* 2017), our results support the idea that needleleaved species primarily adapt to their environment by changing leaf morphology (i.e. SLA) rather than foliar biochemistry (Robakowski *et al.* 2004).

Correlations between leaf nutrient concentrations and traits related to photosynthetic metabolism (*V*_*c,max*_ and *J*_*max*_) are often used to parameterize photosynthesis in ecosystem models (Oleson *et al.* 2013; Rogers *et al.* 2016). We found that these correlations were highly variable between PFTs. Although trait correlations are not necessarily indicative of allocation strategies, this result supports the findings of (Ghimire *et al.* 2017) that N allocation to photosynthesis varies widely by PFT. In tropical evergreen broadleaved trees, for example, photosynthetic metabolism traits were better correlated with *P*_*mass*_ than *N*_*mass*_. This suggests that productivity of tropical species is P-limited (Reich & Oleksyn 2004; Ghimire *et al.* 2017), that N allocation strategies are more variable under N-poor conditions (Ghimire *et al.* 2017), or more generally that photosynthetic metabolism is more sensitive to environmental covariates than leaf nitrogen contents (Ali *et al.* 2015). Meanwhile, the relatively weak *N*_*area*_ *V*_*c,max,area*_ correlation in needleleaved (compared to broadleaved) species echoes earlier results by (Kattge *et al.* 2009) and suggests lower allocation of N to photosynthesis (Ghimire *et al.* 2017). Considering that needleleaf-dominated boreal forests have the largest influence on global climate of any biome (Snyder *et al.* 2004; Bonan 2008), we suggest that parameterization of needleleaf tree productivity based on foliar nitrogen content in Earth System Models be treated with caution.

Correlations of all traits with leaf lifespan were weaker (and often insignificant) within most PFTs than globally. This suggests that leaf economic relationships related to leaf lifespan are dominated by fundamental differences between deciduous and evergreen PFTs, while factors driving variability in leaf lifespan within PFTs are more complex and idiosyncratic (Reich *et al.* 2014; Wu *et al.* 2016a). However, much of this within-PFT variability is driven by variations in shade responses, and a key limitation of our study is the absence of any information about the relative canopy positions at which traits were collected (Lusk *et al.* 2008; Keenan & Niinemets 2016).

Across PFTs, the interaction between growth form and biome in PFT definitions (Table 1) confounds the interpretation of our results with respect to well established biogeographic patterns. We observed as expected that arctic grasses had lower mean SLA than temperate grasses, and that evergreen trees had lower SLA than their deciduous counterparts (Poorter *et al.* 2009). However, by far our highest SLA values were for temperate deciduous broadleaf trees, rather than in grass PFTs as expected (Poorter *et al.* 2009). Similarly to (Onoda *et al.* 2011), we found no consistent patterns in SLA with temperature: Among broadleaved evergreen PFTs, temperate species had lower SLA than tropical, but among broadleaved deciduous PFTs, temperate species had higher SLA than both tropical and boreal species. Unlike (Reich & Oleksyn 2004), who found that foliar N:P ratios decline with latitude, our *N*_*mass*_ estimates were higher in PFTs from colder biomes compared to warmer ones while *P*_*mass*_ was mostly constant between biomes. Contrary to (Atkin *et al.* 2015), our results for both *R*_*d,mass*_ and *R*_*d,area*_ failed to show a trend with respect to biome. However, this comparison may not be entirely fair because our study design inherently averages over the extensive climatic variability within PFTs.

**Table 1:**
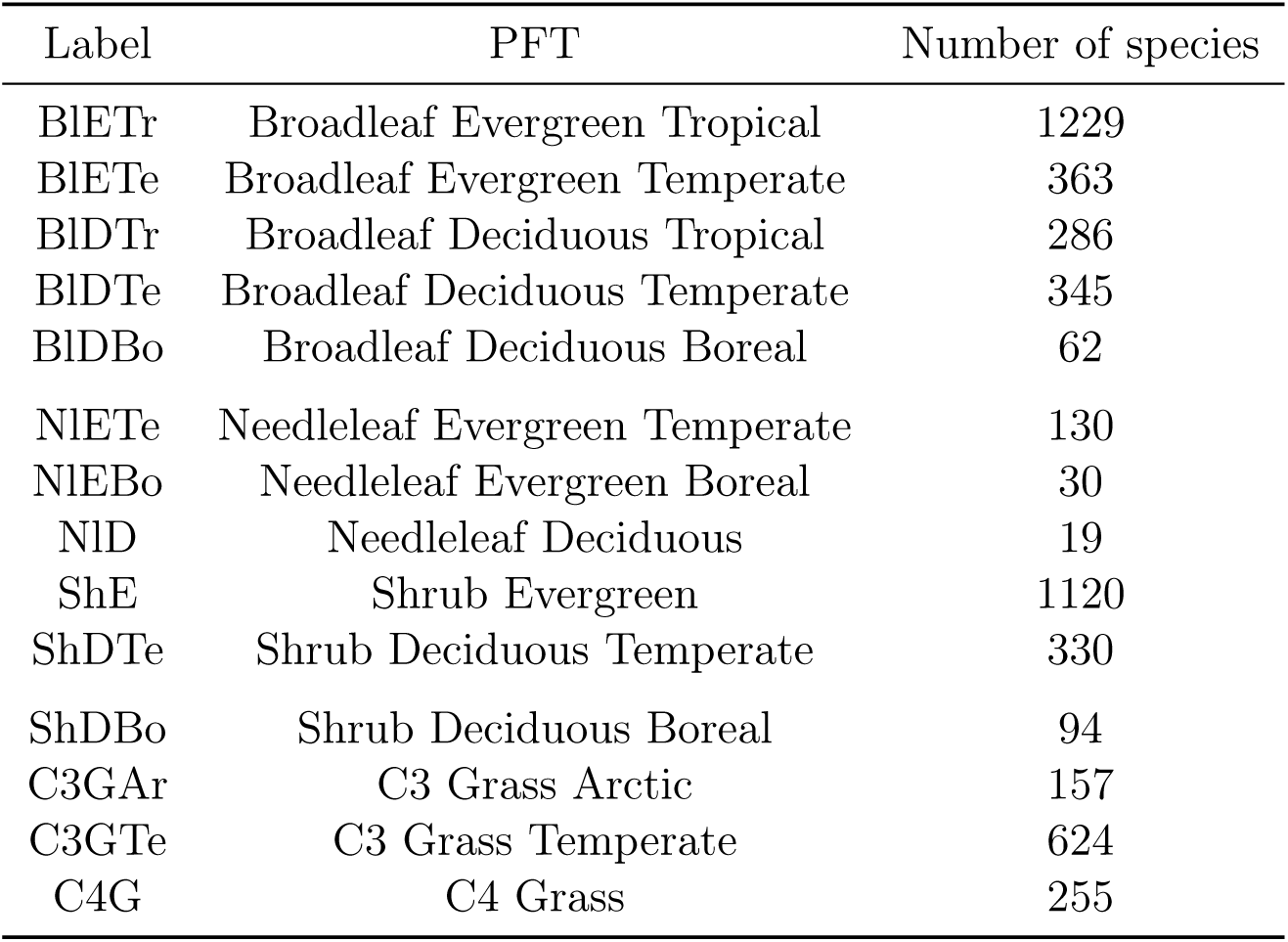
Names, labels, and species counts for plant functional types (PFTs) used in this analysis.

Finally, there has been some debate about the use of mass- or area-normalized traits in analyses of the leaf economic spectrum. Two studies (Lloyd *et al.* 2013; Osnas *et al.* 2013) independently concluded that leaf economic relationships among mass-based traits emerge inevitably out of variation in SLA and are therefore not ecologically meaningful. Responses to these criticisms have suggested that both mass- and area-based normalization have merit: mass-based traits have a natural interpretation in terms of resource allocation, while area-based traits are tied to the area-based nature of energy and gas fluxes through leaf surfaces (Poorter *et al.* 2013; Westoby *et al.* 2013). We argue that investigation of trait correlations on both a mass- and area-basis can yield biologically meaningful conclusions. For one, our discussion of differences in ecological strategies between broadleaved and needleaved species fundamentally depends on comparative analysis of mass- and area-normalized nutrient contents. Meanwhile, our discussion of tropical tree productivity with respect to foliar nutrient contents is supported regardless of how traits are normalized.

### 4.2 Covariance as constraint

The second objective of this paper was to investigate the ability of trait covariance to reduce uncertainties in trait estimates. We show that accounting for covariance reduced uncertainty around PFT-level trait means, particularly for undersampled trait-PFT combinations (Fig. 2 and 3). Moreover, accounting for covariance occasionally changed the *position* of trait mean estimates, even for well-sampled PFT-trait combinations (e.g. *N*_*mass*_ for temperate broadleaved deciduous trees, Fig. 2). This result echoes (Díaz *et al.* 2015) in demonstrating the importance of studying the multivariate trait space rather than individual traits. Such shifts suggest that sampling of these traits in TRY is not representative (Fig. 1; see also Kattge *et al.* 2011). These shifts also indicate that parameter estimates based on univariate trait data (e.g. LeBauer *et al.* 2013; Dietze *et al.* 2014; Butler *et al.* 2017) may not only overestimate uncertainty, but may also be systematically biased. Although some traits in our analysis (R_*d*_, *V*_*c,max*_, and *J*_*max*_) still had insufficient observations necessary to reliably estimate covariance patterns for some PFTs, we show that leveraging covariance increases the effective sample size of all traits. This means that field and remote sensing studies that estimate only certain traits (like SLA and *N*_*mass*_) may be able to use trait correlations to provide constraint on traits they do not directly observe (such as *P*_*mass*_ and *R*_*d,mass*_) (Serbin *et al.* 2014; Musavi *et al.* 2015; Singh *et al.* 2015; Lepine *et al.* 2016). As such, future observational campaigns should consider trait covariance when deciding which traits to measure.

The additional benefit of hierarchical multivariate modeling in our study was limited, largely due to the low number of points used to estimate across-PFT covariance. Therefore, for parameterizing the current generation of ecosystem models using well-sampled traits, simple multivariate models fit independently to each PFT may be sufficient and the additional conceptual challenges and computational overhead of hierarchical modeling are not required. However, for modeling larger numbers of PFTs (Boulangeat *et al.* 2012) and especially individual species (e.g. Linkages Post & Pastor 1996), the benefits of hierarchical modeling may accumulate (Clark 2004; Dietze *et al.* 2008; Cressie *et al.* 2009; Webb *et al.* 2010).

More generally, we foresee tremendous potential for multivariate and hierarchical modeling to elucidate the relationship between traits and organismal and ecosystem function. A natural next step to this study would be to apply the same approach to traits whose relationship to the leaf economic spectrum is less clear. One example is hydraulic traits: While stem and leaf hydraulic traits are correlated (Bartlett *et al.* 2016), the scaling between hydraulic and leaf economic traits is poorly understood (Reich 2014; Li *et al.* 2015). Similarly, reexamining the relationships defining wood (Chave *et al.* 2009; Baraloto *et al.* 2010; Fortunel *et al.* 2012) and root (Kramer-Walter *et al.* 2016; Valverde-Barrantes & Blackwood 2016) economic spectra, as well as their relationship to the foliar traits, would provide useful information on scale-dependence of plant growth and allocation strategies. We emphasize that the difficulty of measuring hydraulic and other non-foliar traits (e.g. Jansen *et al.* 2015) further increases the value of any technique that can fully leverage the information they provide. Ultimately, multivariate and hierarchical modeling may reveal functional trade-offs that are mutually confounding at different scales, thereby enhancing our understanding of processes driving functional diversity.

### 4.3 Conclusions

The vast diversity of plants is a major challenge for functional ecology and ecosystem modeling. Functional diversity research fundamentally depends on dimensionality reduction through a search for meaningful pattern that can be exploited to take reasonable guesses at average behavior. The trait trade-offs comprising the leaf economic spectrum are one such pattern. In this paper, we reaffirm the existence of the leaf economic spectrum both globally and, with some caveats, within plant functional types typically used in the current generation of Earth System Models. We also highlight how the strength of leaf economic relationships can be influenced by biotic and abiotic factors specific to certain PFTs. Finally, we show how patterns of trait covariance like the leaf economic spectrum can be leveraged to inform trait estimates, particularly at small sample sizes.

## Supporting information

## Author contributions

ANS wrote the manuscript and implemented the analysis. ANS and EMC designed the analysis and figures. MCD conceived the original idea for the manuscript, guided its development, and provided financial support. MB, SJ, KK, ÜN, and NAS provided extensive feedback on multiple drafts of the manuscript, and contributed data. CB and YO contributed data.

