## Supplementary material for "Does the leaf economic spectrum hold within plant functional types? A Bayesian multivariate trait meta-analysis"

**Appendix S1:** References for the original sources of the TRY trait data used in our analysis.


Adler, Peter B. et al. (2004). "Functional Traits of Graminoids in Semi-Arid Steppes: a Test of Grazing Histories". In: *Journal of Applied Ecology* 41.4, pp. 653–663. ISSN: 1365-2664. DOI: 10.1111/j.0021-8901.2004.00934.x. URL: https://doi.org/10.1111/j.0021-8901.2004.00934.x.

Adriaenssens, Sandy (2012). *Dry deposition and canopy exchange for temperate tree species under high nitrogen deposition.*

Akhmetzhanova, Asem A. et al. (2012). "A Rediscovered Treasure: Mycorrhizal Intensity Database for 3000 Vascular Plant Species Across the Former Soviet Union". In: *Ecology* 93.3, pp. 689–690. ISSN: 0012-9658. DOI: 10.1890/11-1749.1. URL: https://doi.org/10.1890/11-1749.1.

Araujo, A.C. de et al. (2012). *LBA-ECO CD-02 C and N Isotopes in Leaves and Atmospheric CO2, Amazonas, Brazil.* eng. DOI: 10.3334/ornldaac/1097. URL: https://doi.org/10.3334/ornldaac/1097.

Atkin, Owen K. et al. (1999). "The Response of Fast- and Slow-Growing Acacia Species To Elevated Atmospheric CO2: An Analysis of the Underlying Components of Relative Growth Rate". In: *Oecologia* 120.4, pp. 544–554. ISSN: 1432-1939. DOI: 10.1007/s004420050889. URL: https://doi.org/10.1007/s004420050889.

Auger, Sébastien and Bill Shipley (2012). "Inter-Specific and Intra-Specific Trait Variation Along Short Environmental Gradients in an Old-Growth Temperate Forest". In: *Journal of Vegetation Science* 24.3. Ed. by FrancescoEditor de Bello, pp. 419–428. ISSN: 1100-9233. DOI: 10.1111/j.1654-1103.2012.01473.x. URL: https://doi.org/10.1111/j.1654-1103.2012.01473.x.

Bakker, C., J. Rodenburg, and P. M. van Bodegom (2005). "Effects of Ca- and Fe-Rich Seepage on P Availability and Plant Performance in Calcareous Dune Soils". In: *Plant and Soil* 275.1-2, pp. 111–122. ISSN: 1573-5036. DOI: 10.1007/s11104-005-0438-1. URL: https://doi.org/10.1007/s11104-005-0438-1.

Bakker, C., P. M. Van Bodegom, et al. (2006). "Plant Responses To Rising Water Tables and Nutrient Management in Calcareous Dune Slacks". In: *Plant Ecology* 185.1, pp. 19–28. ISSN: 1573-5052. DOI: 10.1007/s11258-005-9080-5. URL: https://doi.org/10.1007/s11258-005-9080-5.

Baraloto, Christopher et al. (2010). "Decoupled Leaf and Stem Economics in Rain Forest Trees". In: *Ecology Letters* 13.11, pp. 1338–1347. DOI: 10.1111/j.1461-0248.2010.01517.x. URL: https://doi.org/10.1111/j.1461-0248.2010.01517.x.

Beckmann, Michael et al. (2012). "The Role of UV-B Radiation in the Invasion of Hieracium pilosella-A Comparison of German and New Zealand Plants". In: *Environmental and Experimental Botany* 75, pp. 173–180. ISSN: 0098-8472. DOI: 10.1016/j.envexpbot.2011.09.010. URL: https://doi.org/10.1016/j.envexpbot.2011.09.010.

Blonder, Benjamin, Vanessa Buzzard, et al. (2012). "The Leaf-Area Shrinkage Effect Can Bias Paleoclimate and Ecology Research". In: *American Journal of Botany* 99.11, pp. 1756–1763. ISSN: 0002-9122. DOI: 10.3732/ajb.1200062. URL: https://doi.org/10.3732/ajb.1200062.

Blonder, Benjamin, Cyrille Violle, Lisa Patrick Bentley, et al. (2010). "Venation Networks and the Origin of the Leaf Economics Spectrum". In: *Ecology Letters* 14.2, pp. 91–100. ISSN: 1461-023X. DOI: 10.1111/j.1461-0248.2010.01554.x. URL: https://doi.org/10.1111/j.1461-0248.2010.01554.x.

Blonder, Benjamin, Cyrille Violle, and Brian J. Enquist (2013). "Assessing the Causes and Scales of the Leaf Economics Spectrum Using Venation Networks Inpopulus Tremuloides". In: *Journal of Ecology* 101.4. Ed. by HansEditor Cornelissen, pp. 981–989. ISSN: 0022-0477. DOI: 10.1111/1365-2745.12102. URL: https://doi.org/10.1111/1365-2745.12102.

Bocanegra, Kelly, Fernando Fernández, and Jeferson Galvis (2015). “Grupos Funcionales De Árboles En Bosques Secundarios De La Región Bajo Calima (Buenaventura, Colombia)”. In: *Boletín Científico. Centro de Museos. Museo de Historia Natural* 19.1, pp. 17–40. ISSN: 0123-3068. DOI: 10.17151/bccm.2015.19.1.2. URL: https://doi.org/10.17151/bccm.2015.19.1.2.

Bodegom, Peter M. van et al. (2008). “Separating the Effects of Partial Submergence and Soil Oxygen Demand on Plant Physiology”. In: *Ecology* 89.1, pp. 193–204. ISSN: 0012-9658. DOI: 10.1890/07-0390.1. URL: https://doi.org/10.1890/07-0390.1.

Bond-Lamberty, B. et al. (2002). “Leaf Area Dynamics of a Boreal Black Spruce Fire Chronosequence”. In: *Tree Physiology* 22.14, pp. 993–1001. ISSN: 1758-4469. DOI: 10.1093/treephys/22.14.993. URL: https://doi.org/10.1093/treephys/22.14.993.

Bond-Lamberty, B, C Wang, and S T Gower (2002). “Aboveground and Belowground Biomass and Sapwood Area Allometric Equations for Six Boreal Tree Species of Northern Manitoba”. In: *Canadian Journal of Forest Research* 32.8, pp. 1441–1450. ISSN: 1208-6037. DOI: 10.1139/x02-063. URL: https://doi.org/10.1139/x02-063.

Bond-Lamberty, Ben, Stith T. Gower, et al. (2006). “Nitrogen Dynamics of a Boreal Black Spruce Wildfire Chronosequence”. In: *Biogeochemistry* 81.1, pp. 1–16. ISSN: 1573-515X. DOI: 10.1007/s10533-006-9025-7. URL: https://doi.org/10.1007/s10533-006-9025-7.

Bond-Lamberty, Ben, Chuankuan Wang, and Stith T. Gower (2004). “Net Primary Production and Net Ecosystem Production of a Boreal Black Spruce Wildfire Chronosequence”. In: *Global Change Biology* 10.4, pp. 473–487. ISSN: 1365-2486. DOI: 10.1111/j.1529-8817.2003.0742.x. URL: https://doi.org/10.1111/j.1529-8817.2003.0742.x.

Brown, Kerry A. et al. (2011). “Assessing Natural Resource Use By Forest-Reliant Communities in Madagascar Using Functional Diversity and Functional Redundancy Metrics”. In: *PLoS ONE* 6.9. Ed. by NicolasEditor Mouquet, e24107. ISSN: 1932-6203. DOI: 10.1371/journal.pone.0024107. URL: https://doi.org/10.1371/journal.pone.0024107.

Burrascano, S. et al. (2015). “Wild Boar Rooting Intensity Determines Shifts in Understorey Composition and Functional Traits”. In: *Community Ecology* 16.2, pp. 244–253. ISSN: 1588-2756. DOI: 10.1556/168.2015.16.2.12. URL: https://doi.org/10.1556/168.2015.16.2.12.

Butterfield, Bradley J. and John M. Briggs (2010). “Regeneration Niche Differentiates Functional Strategies of Desert Woody Plant Species”. In: *Oecologia* 165.2, pp. 477–487. ISSN: 1432-1939. DOI: 10.1007/s00442-010-1741-y. URL: https://doi.org/10.1007/s00442-010-1741-y.

Byun, Chaeho, Sylvie de Blois, and Jacques Brisson (2012). “Plant Functional Group Identity and Diversity Determine Biotic Resistance To Invasion By an Exotic Grass”. In: *Journal of Ecology* 101.1. Ed. by WillEditor Cornwell, pp. 128–139. ISSN: 0022-0477. DOI: 10.1111/1365-2745.12016. URL: https://doi.org/10.1111/1365-2745.12016.

Campbell, Catherine et al. (2007). “Acclimation of Photosynthesis and Respiration Is Asynchronous in Response To Changes in Temperature Regardless of Plant Functional Group”. In: *New Phytologist* 176.2, pp. 375–389. ISSN: 1469-8137. DOI: 10.1111/j.1469-8137.2007.02183.x. URL: https://doi.org/10.1111/j.1469-8137.2007.02183.x.

Campetella, Giandiego et al. (2011). “Patterns of Plant Trait-Environment Relationships Along a Forest Succession Chronosequence”. In: *Agriculture, Ecosystems & Environment* 145.1, pp. 38–48. ISSN: 0167-8809. DOI: 10.1016/j.agee.2011.06.025. URL: https://doi.org/10.1016/j.agee.2011.06.025.

Carswell, F. E. et al. (2000). “Photosynthetic Capacity in a Central Amazonian Rain Forest”. In: *Tree Physiology* 20.3, pp. 179–186. ISSN: 1758-4469. DOI: 10.1093/treephys/20.3.179. URL: https://doi.org/10.1093/treephys/20.3.179.

Cavender-Bares, Jeannine, Adrienne Keen, and Brianna Miles (2006). “Phylogenetic Structure of Floridian Plant Communities Depends on Taxonomic and Spatial Scale”. In: *Ecology* 87.sp7,

S109–S122. ISSN: 0012-9658. DOI: 10.1890/0012-9658(2006)87[109:psofpc]2.0.co;2. URL: https://doi.org/10.1890/0012-9658(2006)87[109:psofpc]2.0.co;2.

Cerabolini, Bruno E. L. et al. (2010). "Can CSR Classification Be Generally Applied Outside Britain?" In: *Plant Ecology* 210.2, pp. 253–261. ISSN: 1573-5052. DOI: 10.1007/s11258-010-9753-6. URL: https://doi.org/10.1007/s11258-010-9753-6.

Chambers, Jeffrey Q. et al. (2004). "Respiration From a Tropical Forest Ecosystem: Partitioning of Sources and Low Carbon Use Efficiency". In: *Ecological Applications* 14.sp4, pp. 72–88. ISSN: 1051-0761. DOI: 10.1890/01-6012. URL: https://doi.org/10.1890/01-6012.

Chen, Yahan et al. (2011). "Leaf Nitrogen and Phosphorus Concentrations of Woody Plants Differ in Responses To Climate, Soil and Plant Growth Form". In: *Ecography* 36.2, pp. 178–184. ISSN: 0906-7590. DOI: 10.1111/j.1600-0587.2011.06833.x. URL: https://doi.org/10.1111/j.1600-0587.2011.06833.x.

Choat, Brendan et al. (2012). "Global Convergence in the Vulnerability of Forests To Drought". In: *Nature* 491.7426, pp. 752–755. ISSN: 1476-4687. DOI: 10.1038/nature11688. URL: https://doi.org/10.1038/nature11688.

Cornelissen, J. H. C. (1996). "An Experimental Comparison of Leaf Decomposition Rates in a Wide Range of Temperate Plant Species and Types". In: *The Journal of Ecology* 84.4, p. 573. ISSN: 0022-0477. DOI: 10.2307/2261479. URL: https://doi.org/10.2307/2261479.

Cornelissen, J. H. C., P. Castro Diez, and R. Hunt (1996). "Seedling Growth, Allocation and Leaf Attributes in a Wide Range of Woody Plant Species and Types". In: *The Journal of Ecology* 84.5, p. 755. ISSN: 0022-0477. DOI: 10.2307/2261337. URL: https://doi.org/10.2307/2261337.

Cornelissen, J. H. C., H. M. Quested, et al. (2004). "Leaf Digestibility and Litter Decomposability Are Related in a Wide Range of Subarctic Plant Species and Types". In: *Functional Ecology* 18.6, pp. 779–786. ISSN: 1365-2435. DOI: 10.1111/j.0269-8463.2004.00900.x. URL: https://doi.org/10.1111/j.0269-8463.2004.00900.x.

Cornelissen, J.H.C. et al. (2003). "Functional Traits of Woody Plants: Correspondence of Species Rankings Between Field Adults and Laboratory-Grown Seedlings?" In: *Journal of Vegetation Science* 14.3, pp. 311–322. ISSN: 1100-9233. DOI: 10.1111/j.1654-1103.2003.tb02157.x. URL: https://doi.org/10.1111/j.1654-1103.2003.tb02157.x.

Cornwell, William K. et al. (2008). "Plant Species Traits Are the Predominant Control on Litter Decomposition Rates Within Biomes Worldwide". In: *Ecology Letters* 11.10, pp. 1065–1071. ISSN: 1461-0248. DOI: 10.1111/j.1461-0248.2008.01219.x. URL: https://doi.org/10.1111/j.1461-0248.2008.01219.x.

Craine, Joseph M., Andrew J. Elmore, et al. (2009). "Global Patterns of Foliar Nitrogen Isotopes and Their Relationships With Climate, Mycorrhizal Fungi, Foliar Nutrient Concentrations, and Nitrogen Availability". In: *New Phytologist* 183.4, pp. 980–992. ISSN: 1469-8137. DOI: 10.1111/j.1469-8137.2009.02917.x. URL: https://doi.org/10.1111/j.1469-8137.2009.02917.x.

Craine, Joseph M., William G. Lee, et al. (2005). "Environmental Constraints on a Global Relationship Among Leaf and Root Traits of Grasses". In: *Ecology* 86.1, pp. 12–19. ISSN: 0012-9658. DOI: 10.1890/04-1075. URL: https://doi.org/10.1890/04-1075.

Craine, Joseph M., Jesse B. Nippert, et al. (2011). "Functional Consequences of Climate Change-Induced Plant Species Loss in a Tallgrass Prairie". In: *Oecologia* 165.4, pp. 1109–1117. ISSN: 1432-1939. DOI: 10.1007/s00442-011-1938-8. URL: https://doi.org/10.1007/s00442-011-1938-8.

Craine, Joseph M., E. Gene Towne, et al. (2012). "Community Traitscape of Foliar Nitrogen Isotopes Reveals N Availability Patterns in a Tallgrass Prairie". In: *Plant and Soil* 356.1-2, pp. 395–403. ISSN: 1573-5036. DOI: 10.1007/s11104-012-1141-7. URL: https://doi.org/10.1007/s11104-012-1141-7.

Craven, D. et al. (2007). “Between and Within-Site Comparisons of Structural and Physiological Characteristics and Foliar Nutrient Content of 14 Tree Species At a Wet, Fertile Site and a Dry, Infertile Site in Panama”. In: *Forest Ecology and Management* 238.1-3, pp. 335–346. ISSN: 0378-1127. DOI: 10.1016/j.foreco.2006.10.030. URL: https://doi.org/10.1016/j.foreco.2006.10.030.

Demey, Andreas et al. (2013). “Nutrient Input From Hemiparasitic Litter Favors Plant Species With a Fast-Growth Strategy”. In: *Plant and Soil* 371.1-2, pp. 53–66. ISSN: 1573-5036. DOI: 10.1007/s11104-013-1658-4. URL: https://doi.org/10.1007/s11104-013-1658-4.

Diaz, S. et al. (2004). “The Plant Traits That Drive Ecosystems: Evidence From Three Continents”. In: *Journal of Vegetation Science* 15.3, pp. 295–304. ISSN: 1100-9233. DOI: 10.1111/j.1654-1103.2004.tb02266.x. URL: https://doi.org/10.1111/j.1654-1103.2004.tb02266.x.

Domingues, Tomas F., Luiz A. Martinelli, and James R. Ehleringer (2013). “Seasonal Patterns of Leaf-Level Photosynthetic Gas Exchange in an Eastern Amazonian Rain Forest”. In: *Plant Ecology & Diversity* 7.1-2, pp. 189–203. ISSN: 1755-1668. DOI: 10.1080/17550874.2012.748849. URL: https://doi.org/10.1080/17550874.2012.748849.

Domingues, Tomas Ferreira et al. (2010). “Co-Limitation of Photosynthetic Capacity By Nitrogen and Phosphorus in West Africa Woodlands”. In: *Plant, Cell & Environment* 33.6, pp. 959–980. ISSN: 1365-3040. DOI: 10.1111/j.1365-3040.2010.02119.x. URL: https://doi.org/10.1111/j.1365-3040.2010.02119.x.

Fitter, A. H. and H. J. Peat (1994). “The Ecological Flora Database”. In: *The Journal of Ecology* 82.2, p. 415. ISSN: 0022-0477. DOI: 10.2307/2261309. URL: https://doi.org/10.2307/2261309.

Fonseca, Carlos Roberto et al. (2000). “Shifts in Trait-Combinations Along Rainfall and Phosphorus Gradients”. In: *Journal of Ecology* 88.6, pp. 964–977. ISSN: 1365-2745. DOI: 10.1046/j.1365-2745.2000.00506.x. URL: https://doi.org/10.1046/j.1365-2745.2000.00506.x.

Frenette-Dussault, Cédric et al. (2011). “Functional Structure of an Arid Steppe Plant Community Reveals Similarities With Grime’s C-S-R Theory”. In: *Journal of Vegetation Science* 23.2. Ed. by PeterEditor Adler, pp. 208–222. ISSN: 1100-9233. DOI: 10.1111/j.1654-1103.2011.01350.x. URL: https://doi.org/10.1111/j.1654-1103.2011.01350.x.

Fyllas, N. M. et al. (2009). “Basin-Wide Variations in Foliar Properties of Amazonian Forest: Phylogeny, Soils and Climate”. In: *Biogeosciences* 6.11, pp. 2677–2708. ISSN: 1726-4189. DOI: 10.5194/bg-6-2677-2009. URL: https://doi.org/10.5194/bg-6-2677-2009.

Gallagher, Rachael V. and Michelle R. Leishman (2012). “A Global Analysis of Trait Variation and Evolution in Climbing Plants”. In: *Journal of Biogeography* 39.10. Ed. by PaulineEditor Ladiges, pp. 1757–1771. ISSN: 0305-0270. DOI: 10.1111/j.1365-2699.2012.02773.x. URL: https://doi.org/10.1111/j.1365-2699.2012.02773.x.

Garnier, E. et al. (2007). “Assessing the Effects of Land-Use Change on Plant Traits, Communities and Ecosystem Functioning in Grasslands: a Standardized Methodology and Lessons From an Application To 11 European Sites”. In: *Annals of Botany* 99.5, pp. 967–985. ISSN: 1095-8290. DOI: 10.1093/aob/mcl215. URL: https://doi.org/10.1093/aob/mcl215.

Givnish, Thomas J., Rebecca A. Montgomery, and Guillermo Goldstein (2004). “Adaptive Radiation of Photosynthetic Physiology in the Hawaiian Lobeliads: Light Regimes, Static Light Responses, and Whole-Plant Compensation Points”. In: *American Journal of Botany* 91.2, pp. 228–246. ISSN: 0002-9122. DOI: 10.3732/ajb.91.2.228. URL: https://doi.org/10.3732/ajb.91.2.228.

Guerin, G. R., H. Wen, and A. J. Lowe (2012). “Leaf Morphology Shift Linked To Climate Change”. In: *Biology Letters* 8.5, pp. 882–886. ISSN: 1744-957X. DOI: 10.1098/rsbl.2012.0458. URL: https://doi.org/10.1098/rsbl.2012.0458.

Gutiérrez, Alvaro G. and Andreas Huth (2012). “Successional Stages of Primary Temperate Rainforests of Chiloé Island, Chile”. In: *Perspectives in Plant Ecology, Evolution and Systematics*

14.4, pp. 243–256. ISSN: 1433-8319. DOI: 10.1016/j.ppees.2012.01.004. URL: https://doi.org/10.1016/j.ppees.2012.01.004.

Guy, Amanda L., Jenalee M. Mischkolz, and Eric G. Lamb (2013). "Limited Effects of Simulated Acidic Deposition on Seedling Survivorship and Root Morphology of Endemic Plant Taxa of the Athabasca Sand Dunes in Well-Watered Greenhouse Trials". In: *Botany* 91.3, pp. 176–181. ISSN: 1916-2804. DOI: 10.1139/cjb-2012-0162. URL: https://doi.org/10.1139/cjb-2012-0162.

Han, Wenxuan et al. (2005). "Leaf Nitrogen and Phosphorus Stoichiometry Across 753 Terrestrial Plant Species in China". In: *New Phytologist* 168.2, pp. 377–385. ISSN: 1469-8137. DOI: 10.1111/j.1469-8137.2005.01530.x. URL: https://doi.org/10.1111/j.1469-8137.2005.01530.x.

Hickler, T (1999). "Plant Functional Types and community characteristics along environmental gradients on Öland's Great Alvar". MA thesis. Lund, Sweden: University of Lund.

Kattge, Jens et al. (2009). "Quantifying Photosynthetic Capacity and Its Relationship To Leaf Nitrogen Content for Global-Scale Terrestrial Biosphere Models". In: *Global Change Biology* 15.4, pp. 976–991. ISSN: 1365-2486. DOI: 10.1111/j.1365-2486.2008.01744.x. URL: https://doi.org/10.1111/j.1365-2486.2008.01744.x.

Kazakou, E. et al. (2006). "Co-Variations in Litter Decomposition, Leaf Traits and Plant Growth in Species From a Mediterranean Old-Field Succession". In: *Functional Ecology* 20.1, pp. 21–30. ISSN: 1365-2435. DOI: 10.1111/j.1365-2435.2006.01080.x. URL: https://doi.org/10.1111/j.1365-2435.2006.01080.x.

Kerkhoff, Andrew J. et al. (2006). "Phylogenetic and Growth Form Variation in the Scaling of Nitrogen and Phosphorus in the Seed Plants". In: *The American Naturalist* 168.4, E103–E122. ISSN: 1537-5323. DOI: 10.1086/507879. URL: https://doi.org/10.1086/507879.

Kichenin, Emilie et al. (2013). "Contrasting Effects of Plant Inter- and Intraspecific Variation on Community-Level Trait Measures Along an Environmental Gradient". In: *Functional Ecology* 27.5. Ed. by Kaoru Kitajima, pp. 1254–1261. DOI: 10.1111/1365-2435.12116. URL: https://doi.org/10.1111/1365-2435.12116.

Kisel, Yael et al. (2012). "Testing the Link Between Population Genetic Differentiation and Clade Diversification in Costa Rican Orchids". In: *Evolution* 66.10, pp. 3035–3052. ISSN: 0014-3820. DOI: 10.1111/j.1558-5646.2012.01663.x. URL: https://doi.org/10.1111/j.1558-5646.2012.01663.x.

Kleyer, M. et al. (2008). "The LEDA Traitbase: a Database of Life-History Traits of the Northwest European Flora". In: *Journal of Ecology* 96.6, pp. 1266–1274. ISSN: 1365-2745. DOI: 10.1111/j.1365-2745.2008.01430.x. URL: https://doi.org/10.1111/j.1365-2745.2008.01430.x.

Kraft, N. J. B., R. Valencia, and D. D. Ackerly (2008). "Functional Traits and Niche-Based Tree Community Assembly in an Amazonian Forest". In: *Science* 322.5901, pp. 580–582. ISSN: 1095-9203. DOI: 10.1126/science.1160662. URL: https://doi.org/10.1126/science.1160662.

Laughlin, Daniel C. et al. (2011). "Climatic Constraints on Trait-Based Forest Assembly". In: *Journal of Ecology* 99.6, pp. 1489–1499. ISSN: 0022-0477. DOI: 10.1111/j.1365-2745.2011.01885.x. URL: https://doi.org/10.1111/j.1365-2745.2011.01885.x.

Loranger, Jessy et al. (2012). "Predicting Invertebrate Herbivory From Plant Traits: Evidence From 51 Grassland Species in Experimental Monocultures". In: *Ecology* 93.12, pp. 2674–2682. ISSN: 0012-9658. DOI: 10.1890/12-0328.1. URL: https://doi.org/10.1890/12-0328.1.

Louault, F. et al. (2005). "Plant Traits and Functional Types in Response To Reduced Disturbance in a Semi-Natural Grassland". In: *Journal of Vegetation Science* 16.2, pp. 151–160. ISSN: 1100-9233. DOI: 10.1111/j.1654-1103.2005.tb02350.x. URL: https://doi.org/10.1111/j.1654-1103.2005.tb02350.x.

Loveys, B. R. et al. (2003). "Thermal Acclimation of Leaf and Root Respiration: an Investigation Comparing Inherently Fast- and Slow-Growing Plant Species". In: *Global Change Biology* 9.6, pp. 895–910. ISSN: 1365-2486. DOI: 10.1046/j.1365-2486.2003.00611.x. URL: https://doi.org/10.1046/j.1365-2486.2003.00611.x.

Manzoni, Stefano et al. (2013). "Optimization of Stomatal Conductance for Maximum Carbon Gain Under Dynamic Soil Moisture". In: *Advances in Water Resources* 62, pp. 90–105. ISSN: 0309-1708. DOI: 10.1016/j.advwatres.2013.09.020. URL: https://doi.org/10.1016/j.advwatres.2013.09.020.

Medlyn, B. E. et al. (1999). "Effects of Elevated [CO2] on Photosynthesis in European Forest Species: a Meta-Analysis of Model Parameters". In: *Plant, Cell & Environment* 22.12, pp. 1475–1495. ISSN: 0140-7791. DOI: 10.1046/j.1365-3040.1999.00523.x. URL: https://doi.org/10.1046/j.1365-3040.1999.00523.x.

Meir, P. et al. (2002). "Acclimation of Photosynthetic Capacity To Irradiance in Tree Canopies in Relation To Leaf Nitrogen Concentration and Leaf Mass Per Unit Area". In: *Plant, Cell and Environment* 25.3, pp. 343–357. ISSN: 1365-3040. DOI: 10.1046/j.0016-8025.2001.00811.x. URL: https://doi.org/10.1046/j.0016-8025.2001.00811.x.

Meir, Patrick et al. (2007). "Photosynthetic Parameters From Two Contrasting Woody Vegetation Types in West Africa". In: *Plant Ecology* 192.2, pp. 277–287. ISSN: 1573-5052. DOI: 10.1007/s11258-007-9320-y. URL: https://doi.org/10.1007/s11258-007-9320-y.

Messier, Julie, Brian J. McGill, and Martin J. Lechowicz (2010). "How Do Traits Vary Across Ecological Scales? a Case for Trait-Based Ecology". In: *Ecology Letters* 13.7, pp. 838–848. ISSN: 1461-0248. DOI: 10.1111/j.1461-0248.2010.01476.x. URL: https://doi.org/10.1111/j.1461-0248.2010.01476.x.

Meziane, D. and B. Shipley (1999). "Interacting Determinants of Specific Leaf Area in 22 Herbaceous Species: Effects of Irradiance and Nutrient Availability". In: *Plant, Cell & Environment* 22.5, pp. 447–459. ISSN: 1365-3040. DOI: 10.1046/j.1365-3040.1999.00423.x. URL: https://doi.org/10.1046/j.1365-3040.1999.00423.x.

Milla, Rubén and Peter B. Reich (2011). "Multi-Trait Interactions, Not Phylogeny, Fine-Tune Leaf Size Reduction With Increasing Altitude". In: *Annals of Botany* 107.3, pp. 455–465. ISSN: 0305-7364. DOI: 10.1093/aob/mcq261. URL: https://doi.org/10.1093/aob/mcq261.

Minden, Vanessa, Sandra Andratschke, et al. (2012). "Plant Trait-Environment Relationships in Salt Marshes: Deviations From Predictions By Ecological Concepts". In: *Perspectives in Plant Ecology, Evolution and Systematics* 14.3, pp. 183–192. ISSN: 1433-8319. DOI: 10.1016/j.ppees.2012.01.002. URL: https://doi.org/10.1016/j.ppees.2012.01.002.

Minden, Vanessa and Michael Kleyer (2011). "Testing the Effect-Response Framework: Key Response and Effect Traits Determining Above-Ground Biomass of Salt Marshes". In: *Journal of Vegetation Science* 22.3, pp. 387–401. ISSN: 1100-9233. DOI: 10.1111/j.1654-1103.2011.01272.x. URL: https://doi.org/10.1111/j.1654-1103.2011.01272.x.

Müller, Sandra C. et al. (2006). "Plant Functional Types of Woody Species Related To Fire Disturbance in Forest-Grassland Ecotones". In: *Plant Ecology* 189.1, pp. 1–14. ISSN: 1573-5052. DOI: 10.1007/s11258-006-9162-z. URL: https://doi.org/10.1007/s11258-006-9162-z.

Niinemets, Ülo (2001). "Global-Scale Climatic Controls of Leaf Dry Mass Per Area, Density, and Thickness in Trees and Shrubs". In: *Ecology* 82.2, pp. 453–469. ISSN: 0012-9658. DOI: 10.1890/0012-9658(2001)082[0453:gsccol]2.0.co;2. URL: https://doi.org/10.1890/0012-9658(2001)082[0453:gsccol]2.0.co;2.

Ogaya, Romà and Josep Peñuelas (2003). "Comparative Field Study of Quercus Ilex and Phillyrea Latifolia: Photosynthetic Response To Experimental Drought Conditions". In: *Environmental and Experimental Botany* 50.2, pp. 137–148. ISSN: 0098-8472. DOI: 10.1016/s0098-8472(03)00019-4. URL: https://doi.org/10.1016/s0098-8472(03)00019-4.

Onoda, Yusuke et al. (2011). “Global Patterns of Leaf Mechanical Properties”. In: *Ecology Letters* 14.3, pp. 301–312. ISSN: 1461-023X. DOI: 10.1111/j.1461-0248.2010.01582.x. URL: https://doi.org/10.1111/j.1461-0248.2010.01582.x.

Ordoñez, Jenny C. et al. (2010). “Plant Strategies in Relation To Resource Supply in Mesic To Wet Environments: Does Theory Mirror Nature?” In: *The American Naturalist* 175.2, pp. 225–239. ISSN: 1537-5323. DOI: 10.1086/649582. URL: https://doi.org/10.1086/649582.

Pahl, Anna T. et al. (2013). “No Evidence for Local Adaptation in an Invasive Alien Plant: Field and Greenhouse Experiments Tracing a Colonization Sequence”. In: *Annals of Botany* 112.9, pp. 1921–1930. ISSN: 1095-8290. DOI: 10.1093/aob/mct246. URL: https://doi.org/10.1093/aob/mct246.

Peco, Begoña et al. (2005). “The Effect of Grazing Abandonment on Species Composition and Functional Traits: the Case of Dehesa Grasslands”. In: *Basic and Applied Ecology* 6.2, pp. 175–183. ISSN: 1439-1791. DOI: 10.1016/j.baae.2005.01.002. URL: https://doi.org/10.1016/j.baae.2005.01.002.

Penuelas, Josep et al. (2009). “Faster Returns on "Leaf Economics" and Different Biogeochemical Niche in Invasive Compared With Native Plant Species”. In: *Global Change Biology* 16.8, pp. 2171–2185. ISSN: 1365-2486. DOI: 10.1111/j.1365-2486.2009.02054.x. URL: https://doi.org/10.1111/j.1365-2486.2009.02054.x.

Pierce, Simon, Guido Brusa, Matteo Sartori, et al. (2012). “Combined Use of Leaf Size and Economics Traits Allows Direct Comparison of Hydrophyte and Terrestrial Herbaceous Adaptive Strategies”. In: *Annals of Botany* 109.5, pp. 1047–1053. ISSN: 0305-7364. DOI: 10.1093/aob/mcs021. URL: https://doi.org/10.1093/aob/mcs021.

Pierce, Simon, Guido Brusa, Ilda Vagge, et al. (2013). “Allocating Csr Plant Functional Types: the Use of Leaf Economics and Size Traits To Classify Woody and Herbaceous Vascular Plants”. In: *Functional Ecology* 27.4. Ed. by KenEditor Thompson, pp. 1002–1010. ISSN: 0269-8463. DOI: 10.1111/1365-2435.12095. URL: https://doi.org/10.1111/1365-2435.12095.

Pierce, Simon, Alessandra Luzzaro, et al. (2007). “Disturbance Is the Principal $\alpha$-scale Filter Determining Niche Differentiation, Coexistence and Biodiversity in an Alpine Community”. In: *Journal of Ecology* 95.4, pp. 698–706. ISSN: 1365-2745. DOI: 10.1111/j.1365-2745.2007.01242.x. URL: https://doi.org/10.1111/j.1365-2745.2007.01242.x.

Pillar, Valério DePatta and Enio E. Sosinski (2003). “An Improved Method for Searching Plant Functional Types By Numerical Analysis”. In: *Journal of Vegetation Science* 14.3, pp. 323–332. ISSN: 1100-9233. DOI: 10.1111/j.1654-1103.2003.tb02158.x. URL: https://doi.org/10.1111/j.1654-1103.2003.tb02158.x.

Poorter, Hendrik et al. (2009). “Causes and Consequences of Variation in Leaf Mass Per Area (LMA): a Meta-Analysis”. In: *New Phytologist* 182.3, pp. 565–588. ISSN: 0028-646X. DOI: 10.1111/j.1469-8137.2009.02830.x. URL: https://doi.org/10.1111/j.1469-8137.2009.02830.x.

Powers, Jennifer S. and Peter Tiffin (2010). “Plant Functional Type Classifications in Tropical Dry Forests in Costa Rica: Leaf Habit Versus Taxonomic Approaches”. In: *Functional Ecology* 24.4, pp. 927–936. ISSN: 0269-8463. DOI: 10.1111/j.1365-2435.2010.01701.x. URL: https://doi.org/10.1111/j.1365-2435.2010.01701.x.

Prentice, I. Colin et al. (2010). “Evidence of a Universal Scaling Relationship for Leaf CO2 Drawdown Along an Aridity Gradient”. In: *New Phytologist* 190.1, pp. 169–180. ISSN: 0028-646X. DOI: 10.1111/j.1469-8137.2010.03579.x. URL: https://doi.org/10.1111/j.1469-8137.2010.03579.x.

Preston, Katherine A., William K. Cornwell, and Jeanne L. DeNoyer (2006). “Wood Density and Vessel Traits As Distinct Correlates of Ecological Strategy in 51 California Coast Range Angiosperms”. In: *New Phytologist* 170.4, pp. 807–818. ISSN: 1469-8137. DOI: 10.1111/j.1469-8137.2006.01712.x. URL: https://doi.org/10.1111/j.1469-8137.2006.01712.x.

Price, Charles A. and Brian J. Enquist (2007). “Scaling Mass and Morphology in Leaves: an Extension of the Wbe Model”. In: *Ecology* 88.5, pp. 1132–1141. ISSN: 0012-9658. DOI: 10.1890/06-1158. URL: https://doi.org/10.1890/06-1158.

Pyankov, Vladimir I., Alexandra V. Kondratchuk, and Bill Shipley (1999). “Leaf Structure and Specific Leaf Mass: the Alpine Desert Plants of the Eastern Pamirs, Tadjikistan”. In: *New Phytologist* 143.1, pp. 131–142. ISSN: 1469-8137. DOI: 10.1046/j.1469-8137.1999.00435.x. URL: https://doi.org/10.1046/j.1469-8137.1999.00435.x.

Quested, Helen M. et al. (2003). “Decomposition of Sub-Arctic Plants With Differing Nitrogen Economies: a Functional Role for Hemiparasites”. In: *Ecology* 84.12, pp. 3209–3221. ISSN: 0012-9658. DOI: 10.1890/02-0426. URL: https://doi.org/10.1890/02-0426.

Reich, Peter B. et al. (2008). “Scaling of Respiration To Nitrogen in Leaves, Stems and Roots of Higher Land Plants”. In: *Ecology Letters* 11.8, pp. 793–801. ISSN: 1461-0248. DOI: 10.1111/j.1461-0248.2008.01185.x. URL: https://doi.org/10.1111/j.1461-0248.2008.01185.x.

Rüger, Nadja et al. (2009). “Response of Recruitment To Light Availability Across a Tropical Lowland Rain Forest Community”. In: *Journal of Ecology* 97.6, pp. 1360–1368. ISSN: 1365-2745. DOI: 10.1111/j.1365-2745.2009.01552.x. URL: https://doi.org/10.1111/j.1365-2745.2009.01552.x.

— (2011). “Determinants of Mortality Across a Tropical Lowland Rainforest Community”. In: *Oikos* 120.7, pp. 1047–1056. ISSN: 0030-1299. DOI: 10.1111/j.1600-0706.2010.19021.x. URL: https://doi.org/10.1111/j.1600-0706.2010.19021.x.

Sandel, B., J. D. Corbin, and M. Krupa (2011). “Using Plant Functional Traits To Guide Restoration: a Case Study in California Coastal Grassland”. In: *Ecosphere* 2.2, art23. ISSN: 2150-8925. DOI: 10.1890/es10-00175.1. URL: https://doi.org/10.1890/es10-00175.1.

Schererlorenzen, M et al. (2007). “Exploring the Functional Significance of Forest Diversity: a New Long-Term Experiment With Temperate Tree Species (BIOTREE)”. In: *Perspectives in Plant Ecology, Evolution and Systematics* 9.2, pp. 53–70. ISSN: 1433-8319. DOI: 10.1016/j.ppees.2007.08.002. URL: https://doi.org/10.1016/j.ppees.2007.08.002.

Schweingruber, F. and W. Landolt (2005). *The xylem database.*

Shiodera, Satomi, Joeni S. Rahajoe, and Takashi Kohyama (2008). “Variation in Longevity and Traits of Leaves Among Co-Occurring Understorey Plants in a Tropical Montane Forest”. In: *Journal of Tropical Ecology* 24.02, pp. 121–133. ISSN: 1469-7831. DOI: 10.1017/s0266467407004725. URL: https://doi.org/10.1017/s0266467407004725.

Shipley, B. (1995). “Structured Interspecific Determinants of Specific Leaf Area in 34 Species of Herbaceous Angiosperms”. In: *Functional Ecology* 9.2, p. 312. ISSN: 0269-8463. DOI: 10.2307/2390579. URL: https://doi.org/10.2307/2390579.

— (2002). “Trade-Offs Between Net Assimilation Rate and Specific Leaf Area in Determining Relative Growth Rate: Relationship With Daily Irradiance”. In: *Functional Ecology* 16.5, pp. 682–689. ISSN: 1365-2435. DOI: 10.1046/j.1365-2435.2002.00672.x. URL: https://doi.org/10.1046/j.1365-2435.2002.00672.x.

Shipley, Bill and Martin J. Lechowicz (2000). “The Functional Co-Ordination of Leaf Morphology, Nitrogen Concentration, and Gas Exchange In40 Wetland Species”. In: *Écoscience* 7.2, pp. 183–194. ISSN: 2376-7626. DOI: 10.1080/11956860.2000.11682587. URL: https://doi.org/10.1080/11956860.2000.11682587.

Shipley, Bill and Thi-Tam Vu (2002). “Dry Matter Content As a Measure of Dry Matter Concentration in Plants and Their Parts”. In: *New Phytologist* 153.2, pp. 359–364. ISSN: 1469-8137.

DOI: 10.1046/j.0028-646x.2001.00320.x. URL: https://doi.org/10.1046/j.0028-646x.2001.00320.x.

Spasojevic, Marko J. and Katharine N. Suding (2012). "Inferring Community Assembly Mechanisms From Functional Diversity Patterns: the Importance of Multiple Assembly Processes". In: *Journal of Ecology* 100.3, pp. 652–661. ISSN: 0022-0477. DOI: 10.1111/j.1365-2745.2011.01945.x. URL: https://doi.org/10.1111/j.1365-2745.2011.01945.x.

Swaine, E.K. (2007). "Ecological and evolutionary drivers of plant community assembly in a Bornean rain forest". PhD thesis. Aberdeen, Scotland, UK: University of Aberdeen.

Tucker, Sally S., Joseph M. Craine, and Jesse B. Nippert (2011). "Physiological Drought Tolerance and the Structuring of Tallgrass Prairie Assemblages". In: *Ecosphere* 2.4, art48. ISSN: 2150-8925. DOI: 10.1890/es11-00023.1. URL: https://doi.org/10.1890/es11-00023.1.

Veihmeyer, F. J. (1956). "Soil Moisture". In: *Pflanze und Wasser / Water Relations of Plants*, pp. 64–123. DOI: 10.1007/978-3-642-94678-3_8. URL: https://doi.org/10.1007/978-3-642-94678-3_8.

Vergutz, L. et al. (2012). *A Global Database of Carbon and Nutrient Concentrations of Green and Senesced Leaves.* DOI: 10.3334/ORNLDAAC/1106. URL: https://doi.org/10.3334/ORNLDAAC/1106.

Vile, D. (2005). *Significations fonctionnelle et ecologique des traits des especes vegetales: Exemple dans une succession post-cultural Mediterraneenne et generalisations.*

Von Holle, Betsy and Daniel Simberloff (2004). "Testing Fox's Assembly Rule: Does Plant Invasion Depend on Recipient Community Structure?" In: *Oikos* 105.3, pp. 551–563. ISSN: 1600-0706. DOI: 10.1111/j.0030-1299.2004.12597.x. URL: https://doi.org/10.1111/j.0030-1299.2004.12597.x.

Weedon, James T. et al. (2009). "Global Meta-Analysis of Wood Decomposition Rates: a Role for Trait Variation Among Tree Species?" In: *Ecology Letters* 12.1, pp. 45–56. ISSN: 1461-0248. DOI: 10.1111/j.1461-0248.2008.01259.x. URL: https://doi.org/10.1111/j.1461-0248.2008.01259.x.

Williams, M., Y.E. Shimabokuro, and E.B. Rastetter (2012). *LBA-ECO CD-09 Soil and Vegetation Characteristics, Tapajos National Forest, Brazil.* DOI: 10.3334/ORNLDAAC/1104. URL: https://doi.org/10.3334/ORNLDAAC/1104.

Willis, Charles G. et al. (2009). "Phylogenetic Community Structure in Minnesota Oak Savanna Is Influenced By Spatial Extent and Environmental Variation". In: *Ecography*, no–no. ISSN: 1600-0587. DOI: 10.1111/j.1600-0587.2009.05975.x. URL: https://doi.org/10.1111/j.1600-0587.2009.05975.x.

Wilson, K. B., D. D. Baldocchi, and P. J. Hanson (2000). "Spatial and Seasonal Variability of Photosynthetic Parameters and Their Relationship To Leaf Nitrogen in a Deciduous Forest". In: *Tree Physiology* 20.9, pp. 565–578. ISSN: 1758-4469. DOI: 10.1093/treephys/20.9.565. URL: https://doi.org/10.1093/treephys/20.9.565.

Wirth, Christian and Jeremy W. Lichstein (2009). "The Imprint of Species Turnover on Old-Growth Forest Carbon Balances - Insights From a Trait-Based Model of Forest Dynamics". In: *Ecological Studies*, pp. 81–113. ISSN: 0070-8356. DOI: 10.1007/978-3-540-92706-8_5. URL: https://doi.org/10.1007/978-3-540-92706-8_5.

Wohlfahrt, G. et al. (1999). "Inter-Specific Variation of the Biochemical Limitation To Photosynthesis and Related Leaf Traits of 30 Species From Mountain Grassland Ecosystems Under Different Land Use". In: *Plant, Cell and Environment* 22.10, pp. 1281–1296. ISSN: 1365-3040. DOI: 10.1046/j.1365-3040.1999.00479.x. URL: https://doi.org/10.1046/j.1365-3040.1999.00479.x.

Wright, Ian J., David D. Ackerly, et al. (2006). "Relationships Among Ecologically Important Dimensions of Plant Trait Variation in Seven Neotropical Forests". In: *Annals of Botany* 99.5, pp. 1003–1015. ISSN: 1095-8290. DOI: 10.1093/aob/mcl066. URL: https://doi.org/10.1093/aob/mcl066.

Wright, Ian J., Peter B. Reich, et al. (2004). "The Worldwide Leaf Economics Spectrum". In: *Nature* 428.6985, pp. 821–827. ISSN: 1476-4687. DOI: 10.1038/nature02403. URL: https://doi.org/10.1038/nature02403.

Wright, Justin P. and Ariana Sutton-Grier (2012). "Does the Leaf Economic Spectrum Hold Within Local Species Pools Across Varying Environmental Conditions?" In: *Functional Ecology* 26.6. Ed. by CarlyEditor Stevens, pp. 1390–1398. ISSN: 0269-8463. DOI: 10.1111/1365-2435.12001. URL: https://doi.org/10.1111/1365-2435.12001.

Wright, S. Joseph et al. (2010). "Functional Traits and the Growth-Mortality Trade-Off in Tropical Trees". In: *Ecology* 91.12, pp. 3664–3674. ISSN: 0012-9658. DOI: 10.1890/09-2335.1. URL: https://doi.org/10.1890/09-2335.1.

Xu, L. and D. D. Baldocchi (2003). "Seasonal Trends in Photosynthetic Parameters and Stomatal Conductance of Blue Oak (Quercus douglasii) Under Prolonged Summer Drought and High Temperature". In: *Tree Physiology* 23.13, pp. 865–877. ISSN: 1758-4469. DOI: 10.1093/treephys/23.13.865. URL: https://doi.org/10.1093/treephys/23.13.865.

Yguel, Benjamin et al. (2011). "Phytophagy on Phylogenetically Isolated Trees: Why Hosts Should Escape Their Relatives". In: *Ecology Letters* 14.11, pp. 1117–1124. ISSN: 1461-023X. DOI: 10.1111/j.1461-0248.2011.01680.x. URL: https://doi.org/10.1111/j.1461-0248.2011.01680.x.

Zapata-Cuartas, Mauricio, Carlos A. Sierra, and Lauren Alleman (2012). "Probability Distribution of Allometric Coefficients and Bayesian Estimation of Aboveground Tree Biomass". In: *Forest Ecology and Management* 277, pp. 173–179. ISSN: 0378-1127. DOI: 10.1016/j.foreco.2012.04.030. URL: https://doi.org/10.1016/j.foreco.2012.04.030.
